# A critical role for MSR1 in vesicular stomatitis virus infection of the central nervous system

**DOI:** 10.1101/2020.06.17.156703

**Authors:** Duomeng Yang, Tao Lin, Andrew G. Harrison, Tingting Geng, Huadong Wang, Penghua Wang

## Abstract

Macrophage scavenger receptor 1 (MSR1) plays an important role in host defense to bacterial infections, M2 macrophage polarization and lipid homeostasis. However, its physiological function in viral pathogenesis remains poorly defined. Herein, we report that MSR1 facilitates vesicular stomatitis virus (VSV) infection in the spinal cord. Msr1-deficient (*Msr1*^-/-^) mice presented reduced morbidity and mortality following lethal VSV infection, along with normal viremia and antiviral innate immune responses, compared to *Msr1*^+/-^ littermates and wild-type mice. Msr1 expression was selectively upregulated in the spinal cord, which was the predominant target of VSV infection. The viral load in the spinal cord was positively correlated with Msr1 expression level and was reduced in *Msr1*^-/-^ mice. Through its extracellular domain, MSR1 interacted with VSV surface glycoprotein and facilitated its cellular entry. In conclusion, our results demonstrate that MSR1 serves as a cellular entry receptor for VSV and facilitates its infection specifically in the spinal cord.

## Introduction

Macrophage scavenger receptor 1 (MSR1; also known as CD204, SCARA1 and SR-A1), a member of Class A scavenger receptors, performs many functions in homeostasis and immunity, including pathogen clearance, lipid metabolism and macrophage polarization (1). MSR1 recognizes a wide range of ligands including oxidized low-density lipoprotein (oxLDL), endogenous proteins, beta-amyloid, lipoteichoic acid (LTA) and bacterial lipopolysaccharide (LPS) (2, 3). The ligand promiscuity of MSR1 is probably due to its function as a component of other pathogen pattern recognition receptor (PRR) signaling complexes. For instance, MSR1 interacts with receptor tyrosine kinase MER (MERTK) to enable apoptotic cell clearance (4). MSR1 partners with TLR4 to sense LPS that leads to activation of nuclear factor-κB (NF-κB) signaling (5), and therefore contributes to either a pro-or anti-inflammatory responses in Alzheimer’s disease, sepsis, cancer and atherosclerosis conditions (6-8). MSR1 recognizes and internalizes dead cells or debris through interaction with spectrin in macrophages (9). MSR1 also promotes M2 macrophage polarization and Th2 cell responses to bacterial infection by suppressing nuclear translocation of interferon regulatory factor 5 (IRF5) (10). The functions of MSR1 during viral infection may vary with virus species and disease conditions. MSR1 signaling may protect mice against lethal herpes simplex virus infection (11), activate autophagy to control Chikungunya virus infection (12) and limit replication-defective adenovirus type 5-elicited hepatic inflammation and fibrosis by promoting M2 macrophage polarization (13). MSR1 appears to function as a carrier of extracellular dsRNA of hepatitis C virus and present it to endosomal TLR3 and cytoplasmic RIG-I like receptors, resulting in an antiviral type I IFN response (14, 15). However, MSR1 signaling could aggravate the pathogenesis of murine hepatitis virus-induced fulminant hepatitis by enhancing neutrophil NETosis formation-induced complement activation (16).

Vesicular stomatitis virus (VSV) is a negative-sense, single-stranded RNA virus of the *Rhabdoviridae* family. This bullet-shaped virus infects a wide range of cells through its surface glycoprotein (VSV-G) (17, 18). Although VSV infection in humans is rare and very mild, it is common in livestock (e.g. cattle, horses and swine) and may produce significant economic losses, particularly in the southwestern states of the Unites States (19). Because of its broad cell tropism, easily genetic engineering and lack of preexisting human immunity against it, VSV is widely used as a model virus for fundamental research and vaccine development (20, 21). Like rabies virus, VSV is a neurotropic virus that primarily infects animals and occasionally humans (22-24). However, although there are decades of research on the mechanism of VSV neurotropism, the effects of animal model, route of inoculation, and the many signaling pathways that affect VSV neurotropism raise many unanswered questions about the mechanisms involved in neurovirulence. Herein, we report that Msr1 facilitates VSV infection in the spinal cord specifically. Msr1 serves as a cellular receptor that is selectively and highly upregulated in the spinal cord following VSV infection. Msr1-deficiency thus renders mice resistant to lethal VSV infection.

## Results

### Msr1 contributes to VSV pathogenesis in mice

MSR1 plays an important role in host defense to microbial infections (1). However, its physiological function in viral pathogenesis may vary significantly with viral species. We investigate this using VSV, a model RNA virus for the study of innate antiviral immune responses. Msr1 mRNA was expressed in various tissues with the highest in spleen (Fig. 1a). We then infected *Msr1^-/-^* and wild-type (WT, C57BL/6) control mice with a lethal dose of VSV. Intriguingly, ∼90% of the WT mice (n=12) succumbed to VSV infection, but only ∼30% *Msr1^-/-^* mice died (n=12) (Fig. 1b). The overall survival rate of *Msr1^-/-^* mice was much higher than that of WT mice (*P*=0.001, Log-Rank test). In addition, the disease severity of WT mice was continuously increased from day 4 after infection and much greater than *Msr1^-/-^* mice (Fig. 1b). Almost all infected WT mice developed severe symptoms such as hind-limb paralysis as well as decreased mobility (Fig. 1c). To validate the phenotypic results, we produced *Msr1*^-/-^ and *Msr1*^-/+^ littermates and repeated VSV infection. We confirmed that Msr1 protein level in *Msr1^+/-^* was the same as that in WT bone marrow-derived macrophages (BMDM), which express abundant Msr1 (Fig. 1d). Consistently, the survival rate and disease score of *Msr1^+/-^* mice infected with VSV were similar with WT mice. ∼ 80% of infected *Msr1^+/-^* mice (n=6) presented severe hind-limb paralysis and succumbed to lethal VSV infection, while none of *Msr1^-/-^* mice (n=6) showed severe symptoms (Fig. 1e). These data suggest that Msr1 contributes to the pathogenesis of VSV-induced disease.

**Fig. 1.**
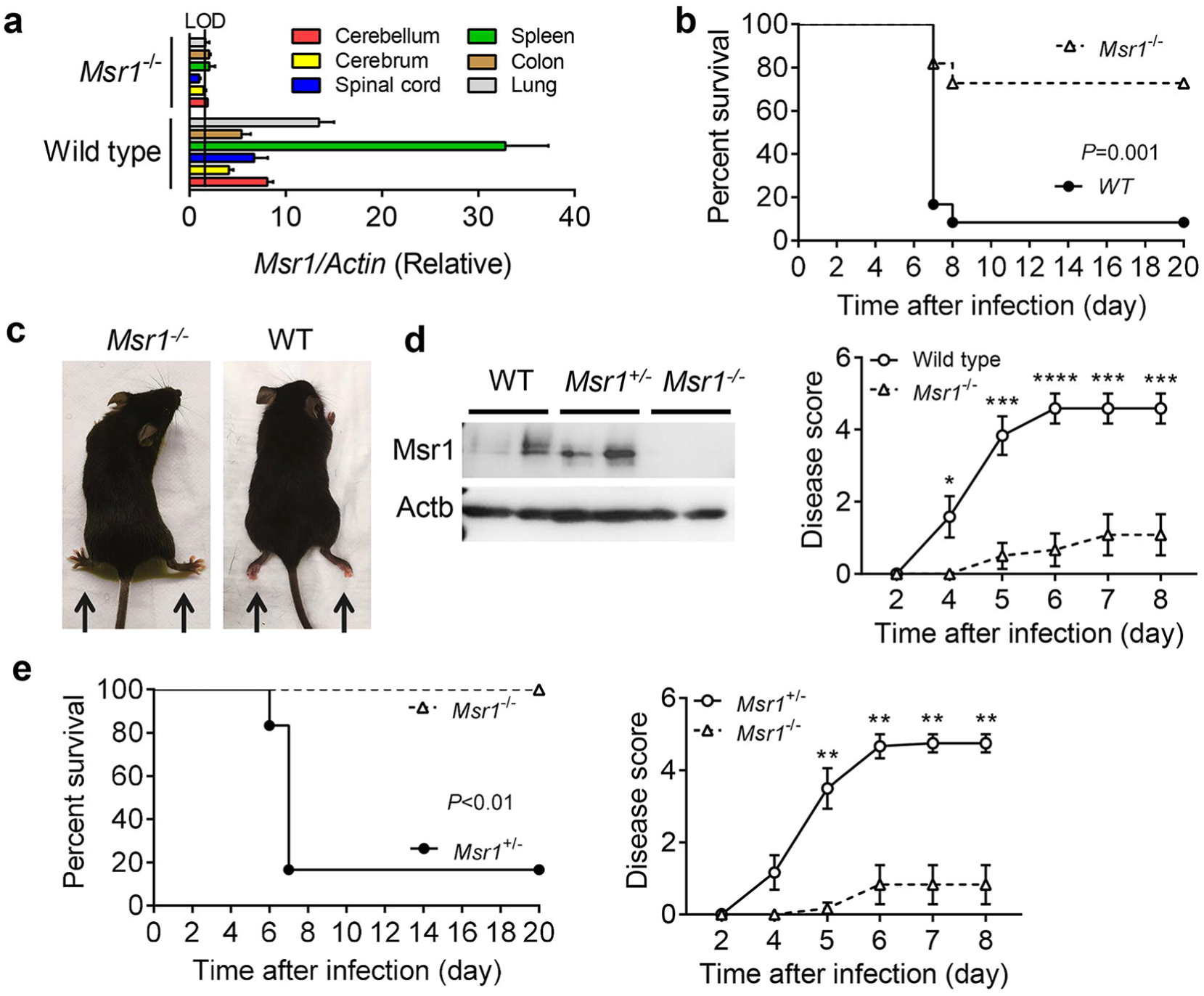
Msr1 contributes to VSV pathogenesis in mice. **a)** *Msr1* mRNA expression in various tissues of wild-type (WT, C57BL/6) and *Msr1^-/-^* mice, N=3 mice per genotype. **b)** The survival curves and disease scores of WT and *Msr1^-/-^* mice challenged with 1×10^7^ plaque-forming units (PFU) per mouse of VSV by retro-orbital injection, N=12 mice/genotype. *P*=0.001 (log-rank test), * *P*<0.05; ** *P*<0.01; *** *P*<0.001; **** *P*<0.0001, non-parametric Mann-Whitney U test. **c)** The WT mice with paralyzed hind-limbs at day 6 post infection. **d)** Immunoblots of Msr1 protein expression in bone marrow-derived macrophages (BMDM) of WT, *Msr1*^-/-^ and *Msr1*^-/+^ littermates. Beta Actin (Actb) is a housekeeping control. **e)** The survival curves and disease scores of *Msr1*^-/-^ and *Msr1*^-/+^ littermates challenged with 1×10^7^ PFU/mouse of VSV by retro-orbital injection, N=6 mice/genotype. All the error bars: mean ± S.E.M.

Msr1 plays an important role in the maintenance of immune homeostasis and is reported to aggravate virus-induced fulminate hepatitis pathogenesis by mediating C5a-induced proinflammatory response(16). We thus assessed VSV viremia and the serum levels of type I IFN and inflammatory cytokines between VSV-infected WT, *Msr1^+/-^* and *Msr1^-/-^* littermates by multiplex ELISA and quantitative RT-PCR. Interestingly, the viremia and cytokine levels (i.e. IFN-α, IFN-γ, TNF-α, IL-10, CXCL-10, IL-1β) in *Msr1^-/-^* mice were similar to WT and *Msr1^+/-^* littermates (Fig. 2a, b, Fig. S1a, b). These results demonstrate that Msr1 is dispensable for systemic VSV dissemination and innate immune responses.

**Fig. 2.**
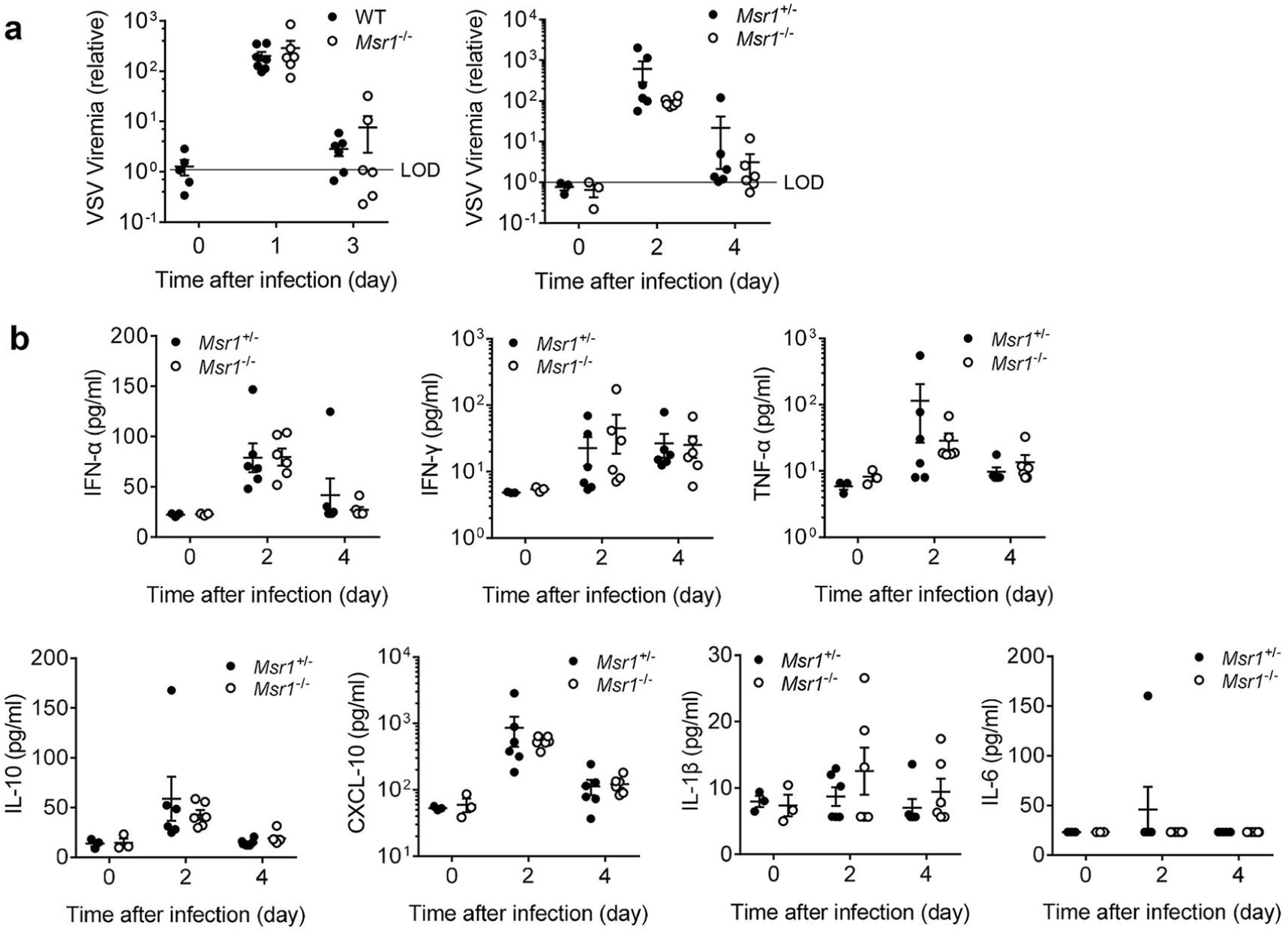
Msr1 is dispensable for systemic VSV dissemination and innate immune responses. **a)** The viral load in whole blood of WT mice, *Msr1^+/-^* and *Msr1^-/-^* littermates infected with VSV, assessed by quantitative RT-PCR, N=6 mice/genotype. **b)** The serum levels of type I IFN and inflammatory cytokines in *Msr1^+/-^* and *Msr1^-/-^* littermates after VSV infection, assessed by multiplex ELISA, N=6 mice/genotype. All the data are presented as mean ± S.E.M. and statistical significance are analyzed by non-parametric Mann-Whitney U test.

### Msr1 is critical for VSV infection in the spinal cord

VSV shows a strong neurotropism and causes severe neurological disorders in a variety of rodent and primate animals though the underlying mechanism remains elusive (25). Indeed, we observed overt hind-limb paralysis and immobility in WT mice (Fig. 1b-e), indicative of severe infection in the central nervous system (CNS). We therefore evaluated viral loads in different tissues after VSV infection. As shown in Fig. 3a, the VSV load in the spinal cord was much higher than other tissues in VSV-infected *Msr1^+/-^* mice. Then, we examined the CNS specimens from VSV-infected *Msr1^+/-^* and *Msr1^-/-^* littermates. We found that the viral load in the spinal cord of *Msr1^-/-^* mice was significantly lower than that of *Msr1^+/-^* mice, while the difference in the brain was moderate (Fig. 3b-d). This result is consistent with VSV-induced severe paraplegia, which is most often caused by spinal cord injury (26). Interestingly, the *Msr1* expression was sharply induced by VSV in the spinal cord and modestly in the liver, while decreased in the spleen after infection (Fig. 3e). A strong positive correlation between *Msr1* expression level and VSV load in the spinal cord was noted; the association was moderate in the brain and spleen (Fig. 3f), suggesting that Msr1 is a determinant of VSV infection in the spinal cord.

**Fig. 3.**
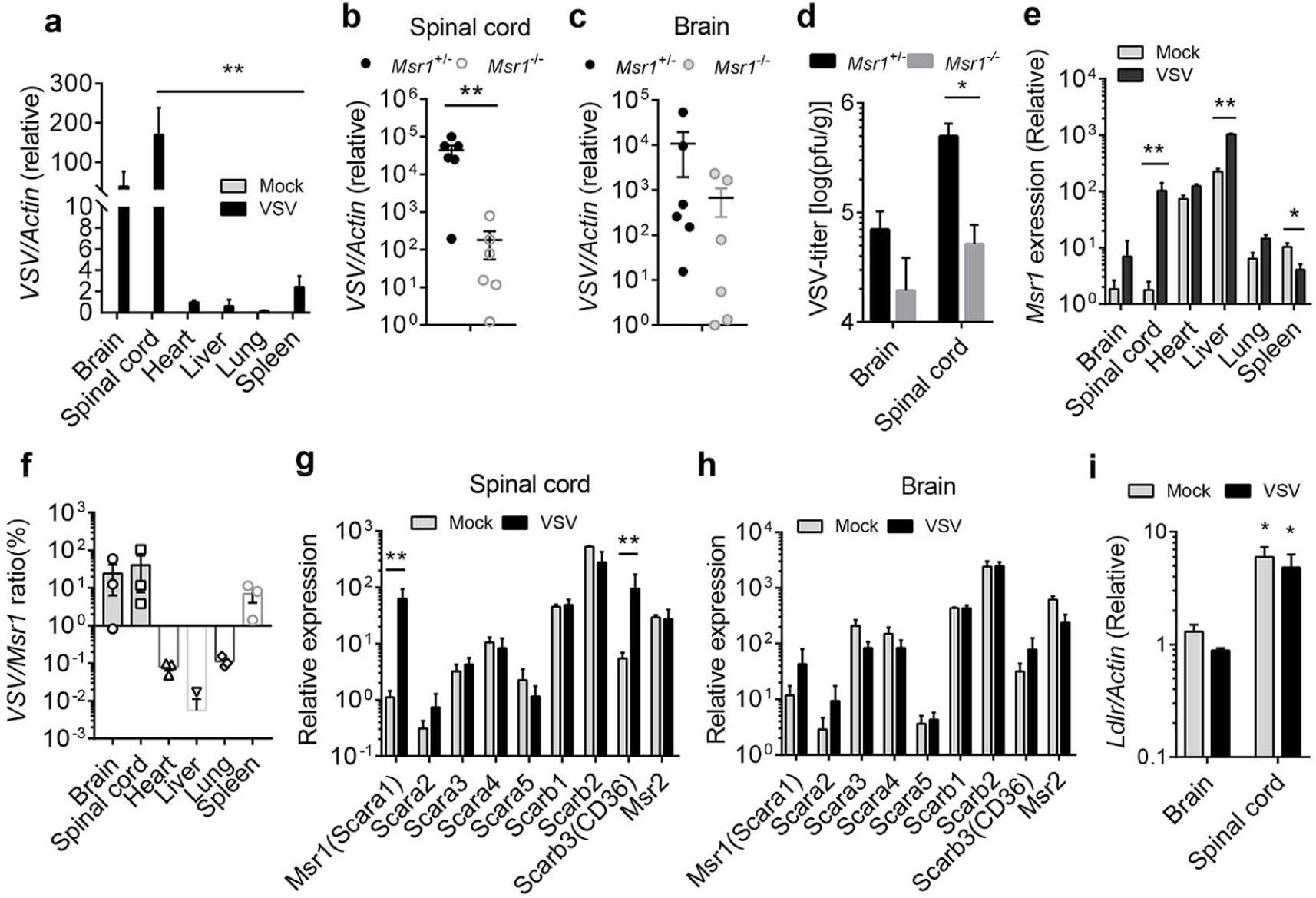
Msr1 is critical for VSV infection specifically in the spinal cord. Quantitative RT-PCR analyses of VSV loads in **a)** different tissues of *Msr1^+/-^* mice (N=3 mice per group), **b)** the spinal cords and **c)** the brains of *Msr1^+/-^* and *Msr1^-/-^* littermates on day 6 after infection (N=6 mice/genotype). **d)** The VSV titers in the spinal cord and brain of *Msr1^+/-^* and *Msr1^-/-^* littermates on day 6 after infection, as assessed by plaque forming assay. N=6 mice/genotype. **e)** Msr1 mRNA expression in different tissues of mock and VSV-infected mice on day 6 after infection, N=3 mice per group. **f)** Correlation between *Msr1* expression and VSV load in various tissues after VSV infection, N=3 mice per group. The mRNA expression of Class A and B scavenger receptors in **g)** the spinal cord and **h)** the brain on day 6 after VSV infection, as assessed by quantitative RT-PCR, N=3 mice per group. **i)** *Ldlr* mRNA expression in the brain and spinal cord, N=3 mice per group. Mock: no virus. All the data are presented as mean ± S.E.M. and statistical significances are analyzed by non-parametric Mann-Whitney U test, * *P*<0.05, ** *P*<0.01.

The scavenger receptor family consists of 10 classes, which are structurally heterogeneous with little or no homology (1). MSR1 belongs to the Class A receptors, which all have a extracellular collagenous domain (1). The aforementioned results suggest that MSR1 is upregulated by VSV specifically in the spinal cord. We then asked if other members of Class A and B may be induced similarly by VSV. The results show that *CD36* (*Scarb3*) was also markedly induced in the spinal cord after VSV infection (Fig. 3g), but none of the Class A/B scavenger receptors tested were significantly changed in the expression levels in the infected brain (Fig. 3h). The LDL receptor family members serve as the cellular receptors for VSV (27), and are also expressed in the nervous system (28); we thus examined the expression of *Ldlr* in the brain and spinal cord. Intriguingly, *Ldlr* expression was higher in the spinal cord than brain, but not significantly changed after VSV infection in either tissue (Fig. 3i). These findings suggest that Msr1 is critical for VSV infection, specifically in the spinal cord.

### Msr1 mediates cellular entry of VSV

The above-mentioned data suggest that MSR1 may facilitate VSV infection directly. Of note, MSR1 is known to recognize a wide range of self-and microbial ligands, in particular, oxidized low density lipoproteins(2, 3). We thus hypothesized that MSR1 could mediate cellular entry of VSV. To this end, we employed primary neuron culturing from neonatal *Msr1^+/-^* and *Msr1^-/-^* littermates and BMDM which express abundant Msr1 (Fig. 4c). Indeed, the VSV load in *Msr1^-/-^* neurons (Fig. 4a) and the virus titer in its culture medium (Fig. 4b) were significantly decreased compared to *Msr1^+/-^* littermates. The intracellular VSV-G protein level and extracellular infectious virions in *Msr1^-/-^* BMDMs were much lower than those in WT cells (Fig. 4d-f). We next determined if cellular entry of VSV was impaired in *Msr1^-/-^* cells. BMDMs were inoculated with VSV at 4 °C for 2hrs to allow for VSV attachment to the cell surface, at 37 °C for 30 min to allow for VSV entry into cells, and lastly were washed extensively to remove unbound virions. Indeed, there were fewer VSV virions attached to *Msr1^-/-^* than WT cells at 4 °C and consequently fewer virions inside *Msr1^-/-^* cells at 37 °C (Fig. 4g). These data show that Msr1 mediates cellular entry of VSV.

**Fig. 4.**
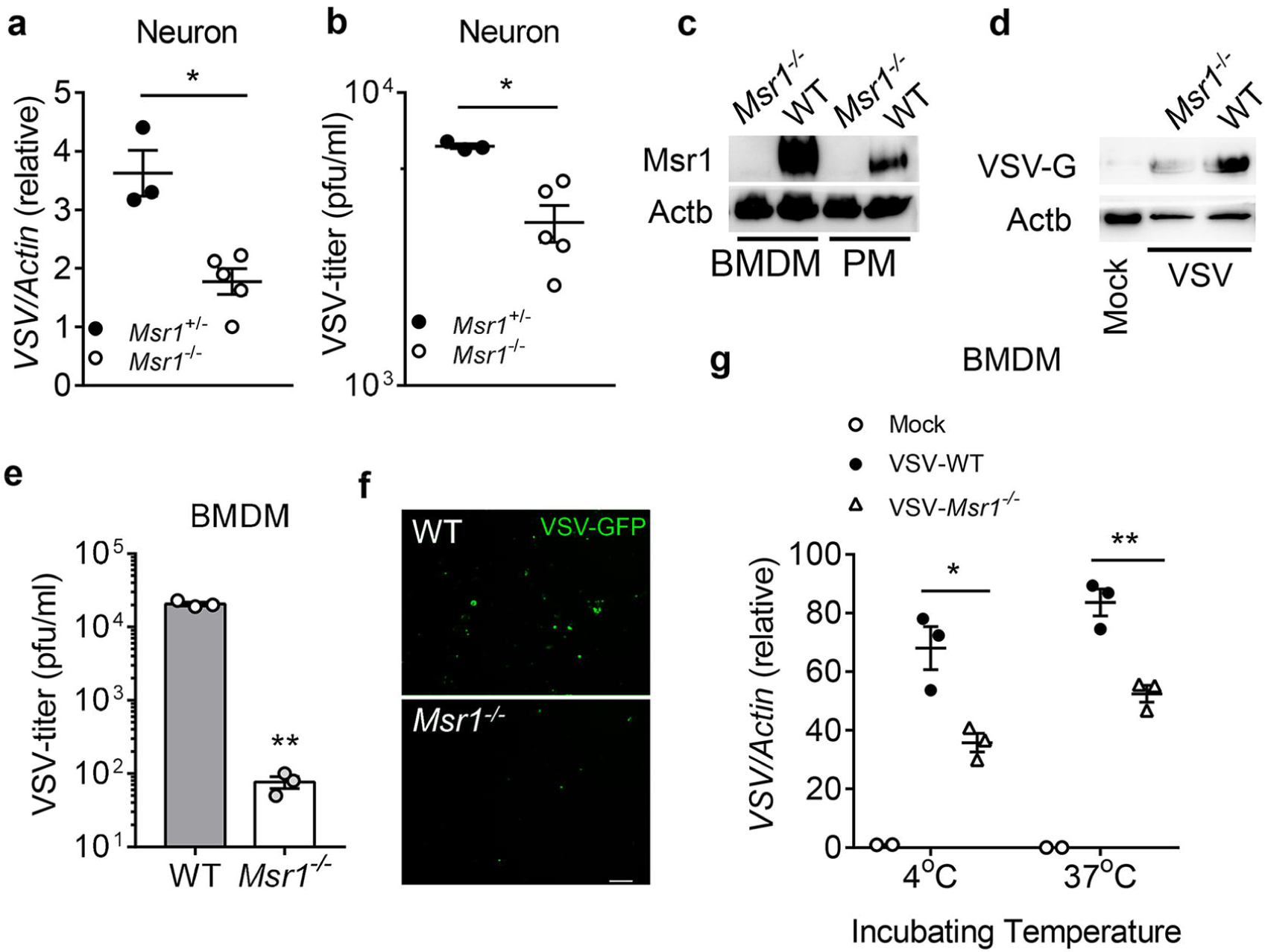
Msr1 mediates cellular entry of VSV in primary mouse cells. VSV loads in primary neurons of *Msr1^+/-^* and *Msr1^-/-^* littermates at 36 h after inoculation (multiplicity of infection =3), as assessed by **a)** quantitative RT-PCR and **b)** plaque forming assay. Each dot represents an individual mouse. **c)** Immunoblots of Msr1 protein expression in WT and *Msr1^-/-^* bone marrow-derived macrophages (BMDM) and peritoneal macrophages (PM). **d)** VSV-G protein expression in BMDM, **e)** virus titer in culture medium and **f)** VSV-GFP fluorescence under microscopy in BMDM at 48 h after VSV infection, N=3 biological replicates. Objective: 20x, Scale bar: 100 µm. **g)** Quantitative RT-PCR analyses of VSV virions attached to BMDM (4°C for 2 h) and entry into BMDM (37°C for 30 min), N=3 biological replicates. Beta Actin (Actb) is a housekeeping control. Mock: no virus. All the data are presented as mean ± S.E.M. and statistical significances are analyzed by a standard two-tailed unpaired Student’s t-test, * *P*<0.05, ** *P*<0.01.

Human MSR1 protein shares 70% identity and 81% similarity with mouse Msr1. We then asked if the function of MSR1during VSV infection is evolutionarily conserved. To this end, we evaluated VSV infection in a human *MSR1^-/-^* trophoblast line that was generated recently in our hands (12). Indeed, VSV-G protein level in *MSR1^-/-^* trophoblasts was much less than that in WT cells at 12 hrs after VSV infection (Fig. 5a). Consistently, the fluorescence intensity of VSV-GFP in *MSR1^-/-^* trophoblasts was also decreased compared to WT cells (Fig. 5b), and the plaque assay further showed a significant reduction in VSV virions in the culture medium of *MSR1^-/-^* compared to WT trophoblasts (Fig. 5c). Overexpression of MSR1 significantly increased VSV-G protein level (Fig. 5d), VSV-GFP fluorescence intensity (Fig. 5e), and VSV titer in the cell culture medium (Fig. 5f), compared to the vector control. Epichromosomal complementation of MSR1 gene in *MSR1^-/-^* trophoblasts restored VSV infection (Fig. 5g).

**Fig. 5.**
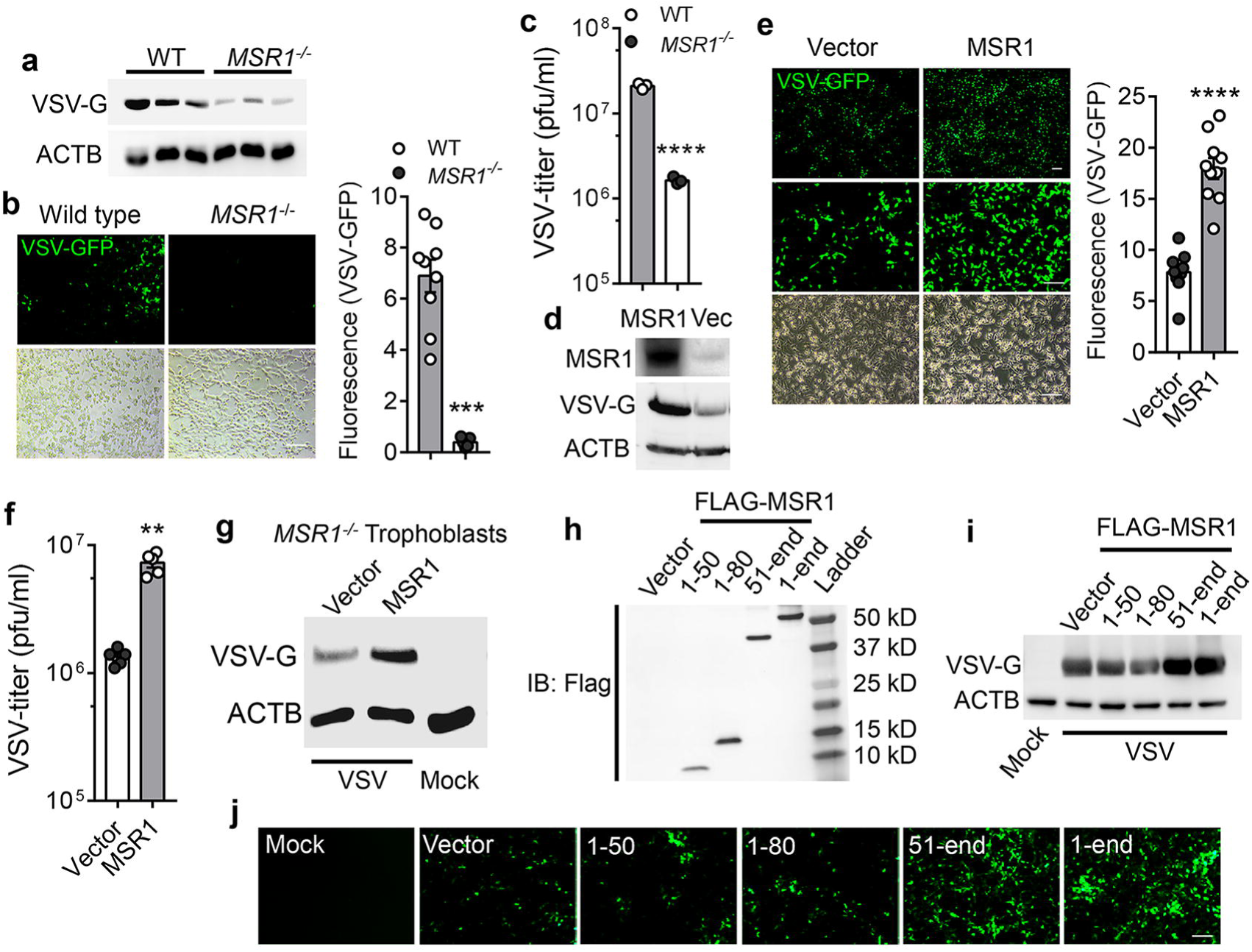
MSR1 facilitates cellular entry of VSV through its extracellular domain. **a)-c)** WT and *MSR1^-/-^* trophoblasts were examined at 12 h post VSV-GFP inoculation, **a)** Immunoblots of VSV-G protein level, **b)** VSV-GFP fluorescence intensity and **c)** VSV titers in the cell culture medium, MOI=0.5. N=3 biological replicates. **d)-f)** WT and MSR1-overexpressed trophoblasts were examined at 12 h post VSV-GFP inoculation, **d)** MSR1 protein expression, VSV-G protein level, **e)** VSV-GFP fluorescence intensity and **f)** VSV titers in the culture medium, MOI=0.5. N=3 biological replicates. The VSV-GFP fluorescence intensity in **b)** and **e)** was calculated from 9 random fields of view from three biological replicates. Objective: 4x (top) and 20x, Scale bar: 100 µm. **g)** Immunoblots of VSV-G protein level in *MSR1^-/-^* trophoblasts with epichromosomal complementation of an empty vector or FLAG-MSR1 expression plasmid at 12 h after VSV infection, MOI=0.5. **h)** The protein expression of different FLAG-tagged MSR1 fragments at 24 h after transfection of plasmids in trophoblasts. IB: immunoblotting. Amino residues 1-50 (cytoplasmic N-tail), 1-80 (cytoplasmic N-tail plus transmembrane), 51-end (extracellular domains with transmembrane), and 1-end (full-length). **i)** VSV-G protein level and **j)** VSV-GFP fluorescence intensity in the trophoblasts overexpressing FLAG-MSR1 fragments at 12 h after VSV-GFP infection, MOI=0.5, N=3 biological replicates. Beta Actin (ACTB) is a housekeeping control. Mock: no virus. All the data are presented as mean ± S.E.M. and statistical significances are analyzed by a standard two-tailed unpaired Student’s t-test, ** *P*<0.01, *** *P*<0.001, **** *P*<0.0001.

MSR1 is a cell surface protein with a short intracellular N-terminus, a transmembrane domain and a long extracellular portion. We postulated that the extracellular portion mediates VSV entry. To prove this, we dissected MSR1 into several fragments: amino residues 1-50 (cytoplasmic N-tail), 1-80 (cytoplasmic N-tail plus transmembrane), and 51-end (extracellular domains with transmembrane), expressed them in trophoblasts (Fig. 5h, Fig. S2a) and infected cells with VSV-GFP for 12 hrs. Intriguingly, both the fragment (51-end) and full-length (1-end) MSR1 enhanced VSV-G protein level and VSV-GFP fluorescence intensity; while the fragments without the extracellular domains (1-50 and 1-80) failed to do so (Fig. 5i, j). These results demonstrate that MSR1 facilitates cellular entry of VSV through its extracellular domain.

### MSR1 interacts with VSV glycoprotein G via its extracellular domain

Based on the aforementioned data, we hypothesized that MSR1 serves as a cellular receptor for VSV. We thus employed an immunoprecipitation assay to investigate if MSR1 full-length (1-end) and/or fragments (1-80, 51-end, respectively) directly interact with intact VSV-GFP virions. In agreement with Fig. 5h-j, both the 51-end and full-length of MSR1 pulled-down infectious virions; while the MSR1 (1-80) did not (Fig. 6a-c). Of note, with considerably less protein expression, the 51-end fragment pulled down the same amount of VSV-GFP as full-length MSR1 (Fig. 6a-c), suggesting that the extracellular fragment mediates MSR1 binding to VSV virions. We further asked if MSR1 directly interacts with VSV surface glycoprotein G that mediates VSV entry into host cells. We performed co-transfection of VSV-G and FLAG-MSR1/fragment plasmids into HEK293 cells and immunoprecipitation using an anti-FLAG antibody. The results show that the fragment (51-end) and full-length (1-end) FLAG-MSR1 pulled down VSV-G, while the empty vector and the fragment 1-80 did not (Fig. 6d).

**Fig. 6.**
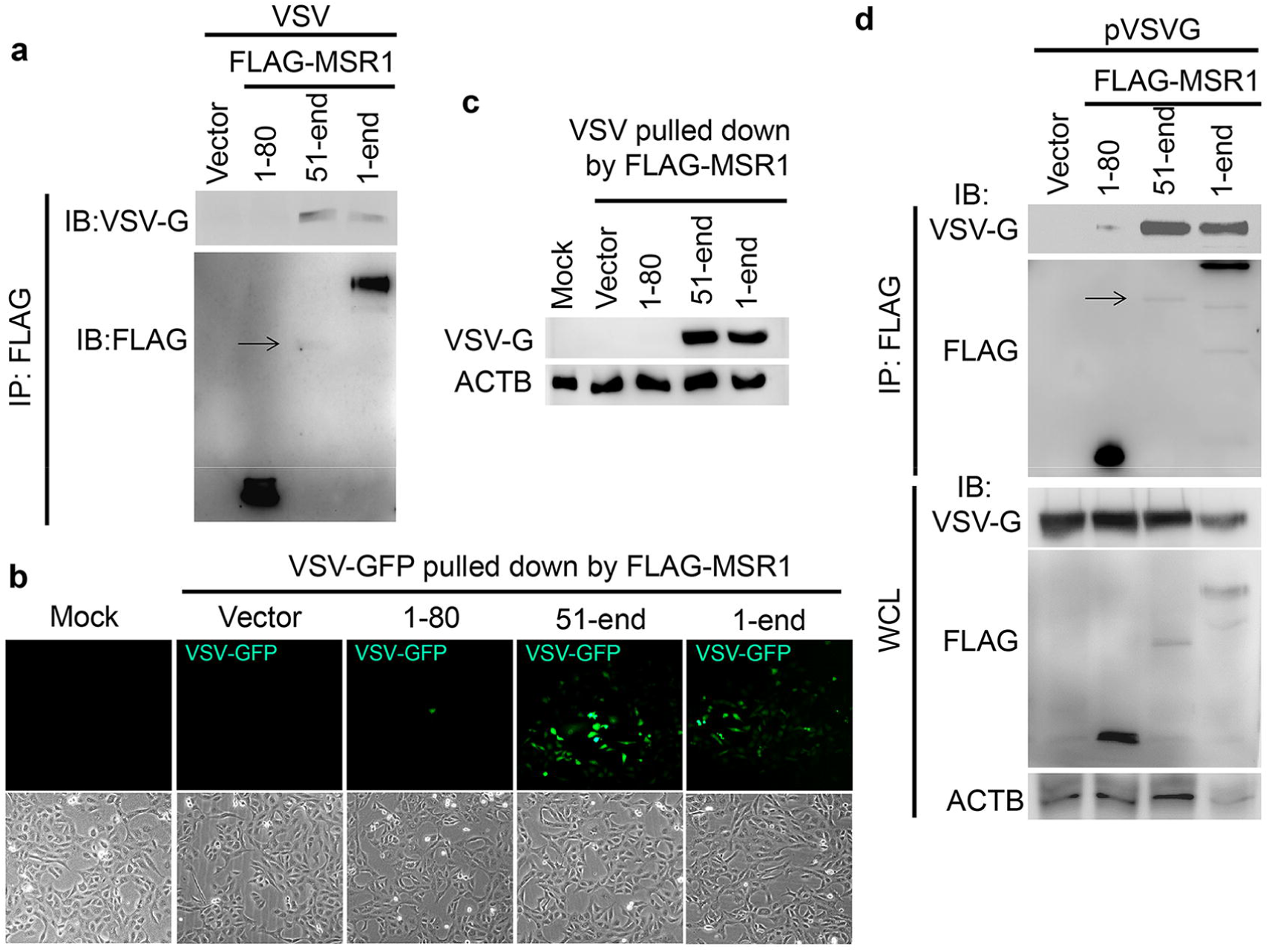
MSR1 interacts with VSV glycoprotein G via its extracellular domains. **a-c)** Binding of MSR1 to VSV virions. FLAG-MSR1 (human, 1-end) and its fragments (aa1-80, 51-end) were expressed in HEK293 cells and immunoprecipitated (IP) using anti-FLAG antibody-coated magnetic beads, which were then incubated with intact VSV-GFP virions (4°C for 2 h). The bound virions were eluted for **a)** immunoblotting (IB) with a mouse monoclonal anti-FLAG and rabbit anti-VSV-G antibody, **b-c**) for re-infection in Vero cells for 12 h followed by detection of **b)** VSV-GFP by fluorescence microscopy and **c)** VSV-G protein level by immunoblotting. Mock: no virion. **d)** co-IP of FLAG-MSR1 and its fragments with VSV-G in HEK293 cells transfected with expression plasmids. WCL: whole cell lysate. Arrows point to the faint protein bands.

## Discussion

Although MSR1 has proven to perform a variety of functions in homeostasis and host immunity due to its ability to recognize a diverse range of ligands (2, 3), its role in viral pathogenesis remains poorly defined. MSR1 signaling may be protective against Chikungunya virus and herpes simplex virus infection (11, 12), but could also contribute to hepatitis virus-induced liver damage (16). In this study, our data collectively demonstrate that MSR1 serves as a cellular receptor facilitating VSV infection specifically in the spinal cord, and this is substantiated by several lines of experimental evidence. Firstly, Msr1-deficient mice were strikingly resistant to VSV-elicited hind-limb paralysis compared to their wild-type littermates. However, Msr1 was dispensable for systemic VSV dissemination and innate immune responses. Secondly, the viral load in the spinal cord was the highest and the fold induction of Msr1 expression by VSV was the greatest among all tested tissues. Of note, the Msr1 expression level was positively correlated with VSV load in the spinal cord. These results not only demonstrate a strong tropism of VSV for the spinal cord that leads to severe paraplegia(26),(25), but also reveal a novel function for Msr1 as a critical determinant of VSV infection specifically in the spinal cord. Thirdly, VSV attachment, entry, and consequently replication was significantly reduced in both mouse primary *Msr1*^-/-^ macrophages and human *MSR1*^-/-^ trophoblasts. While overexpression of MSR1 could enhance cellular entry of VSV and subsequent replication. Lastly, MSR1, through its extracellular domain, interacted with VSV virions through its surface glycoprotein G and overexpression of the extracellular section alone was sufficient to promote VSV infection. These observations suggest that MSR1 is an entry receptor for VSV (17, 18).

To date, the membrane lipid phosphatidylserine and endoplasmic reticulum chaperone Gp96 are important for VSV entry, but not as a receptor (25). The LDL receptor and its family members were recently characterized as entry receptors for VSV in both human and mouse cells (27). The widespread expression pattern of the LDLR family members may thus in part underlie VSV pantropism. However, the *Rhabdoviridae* viruses (e.g. rabies virus, VSV) specifically invade and replicate in motor neurons that exist predominately in the spinal cord (29), and subsequent retrograde or anterograde spread of progeny virions in the motor nerve system results in paraplegia (30, 31). Indeed, this study demonstrates that VSV primarily targets the spinal cord, and secondarily the brain. Intriguingly MSR1 is essential for VSV infection only in the spinal cord. In agreement with this, among all tested Class A and B scavenger receptors, MSR1 is the most significantly upregulated member in the spinal cord; while LDLR expression is unchanged following VSV infection. These observations suggest that MSR1 becomes important for VSV infection of the spinal cord at least partly because MSR1 expression is robustly upregulated. On the other hand, MSR1 could contribute to neuronal inflammation and apoptosis by activating NF-κB signaling pathway in the spinal cord (32). However, our data do not necessarily rule out the importance of LDLR family to VSV infection in the spinal cord. Indeed, LDLR expression in the spinal cord, either before or after VSV infection, is much higher than the brain.

In sum, we herein define MSR1 as a cellular entry receptor for VSV infection specifically in the spinal cord. MSR1 can be selectively and highly upregulated in the spinal cord following VSV infection. Given that VSV and its pseudotyped viruses have been widely applied to vaccine research and oncolytic therapy (33-35), it seems plausible to improve VSV-mediated oncolytic efficacy by manipulating MSR1 expression.

## Materials and Methods

### Animals

All mice were purchased from the Jackson Laboratory and housed under the same conditions. C57BL/6J mice (Stock No: 000664) were employed as the wild-type control mice. *Msr1^-/-^* mice (B6.Cg-*Msr1^tm1Csk^*/J, stock No: 006096) were made from 129 embryonic stem (ES) cells and then backcrossed to C57BL/6J mice for 12 generations. *Msr1^+/-^* mice were made by backcrossing *Msr1^-/-^* mice to C57BL/6J mice. All of the gene-mutant and wild type mice were normal in body weight and general health. Sex-and age-matched (6-8 weeks) mice were used for all the experiments. The animal protocols were approved by the Institutional Animal Care & Use Committee at University of Connecticut adhering to the National institutes of Health recommendations for the care and use of laboratory animals.

### Antibodies and plasmids

The rabbit anti-Actin (Cat # 8456) antibody was purchased from Cell Signaling Technology (Danvers, MA 01923, USA). The mouse anti-FLAG (Cat# TA50011) and rabbit anti-human/mouse MSR1 (Cat# TA336699) antibodies were from Origene (Rockville, MD 20850, USA). The mouse anti-human MSR1 (Cat# MAB2708) antibody was obtained from R&D Systems (Minneapolis, MN 55413, USA). Then rabbit anti-VSV-G (Cat#V4888) antibody and the anti-FLAG M2 magnetic beads (Clone M2, Cat# M8823) were from Sigma-Aldrich (St. Louis, MO, USA). The human MSR1 (Cat# RC209609) were obtained from Origene (Rockville, MD 20850, USA). Human MSR1 and its fragment 1-50, 1-80 and 51-end were amplified by PCR and inserted into pcDNA3.1 (Zeo)-FLAG vector. The primers are listed in the Supplemental Table S1.

### Cell culture and viruses

Vero cells (monkey kidney epithelial cells, # CCL-81), human trophoblasts HTR-8/SVneo (#CRL-3271) and L929 (mouse fibroblast cells, # CCL-1) were purchased from American Type Culture Collection (ATCC) (Manassas, VA20110, USA). *MSR1^-/-^* trophoblasts were generated in our previous study (12). Vero cells were grown in Corning® DMEM (Dulbecco’s Modified Eagle’s Medium) supplemented with 10% FBS and 1% penicillin/streptomycin. Human trophoblasts were cultured in Corning® RPMI 1640 medium supplemented with 10% fetal bovine serum and 1% penicillin/streptomycin. Bone marrow-derived macrophages (BMDMs) were differentiated from femur bone-derived bone marrows in L929 conditioned medium (RPMI1640, 20%FBS, 30% L929 culture medium and 1% penicillin/streptomycin) in 10-cm Petri-dishes at 37 °C, 5% CO_2_ for 7 days. The BMDMs were then seeded in culture plates and cultured in regular RPMI1640 medium overnight before further treatment. MycoZap (Lonza) were used regularly to prevent mycoplasma contamination. The vesicular stomatitis virus (VSV) used in this study was Indiana strain described previously (36), and the green fluorescence protein (GFP) tagged VSV was made from this VSV. These viruses were propagated in Vero cells.

### Primary neuron culture

The primary neuron culturing was performed according to a previous method(37). The brain of a 3-day old neonate was dissected dorsally under a microscope; the vascular tunic was removed; and then the brain was washed in ice-cold DMEM medium. The brain was then broken into pieces by pipetting up and down in 0.125% trypsin with DNAse I, and then digested for 30min in a 37 °C water bath, vortexed every 10 min. Digestion of the brain was terminated by 10% FBS+DMEM, filtered through a 70μm cell strainer (ThemoFisher brand). The filtrate was centrifuged at 500 x g at 4 °C for 5min to pellet neurons. The neurons were then plated in 0.0125% poly-L-lysine-coated culture plates (overnight at 37°C), and cultured in neurobasal medium (Gibco, U.S) with 2% B27 supplement (Gibco, U.S) at 37 °C, 5% CO_2_ for 5-6 days.

### Mouse infection and disease monitoring

VSV stock [∼10^8^ plaque forming units (PFU/ml)] was diluted in sterile phosphate buffered saline (PBS), and ∼10^7^ PFU was injected into mice retro-orbitally (38). The mice were monitored for disease progression over a period of 8 days after infection. The main symptoms were mobility and hind-limb paralysis. The severity of disease was measured using a scale from 0 (no symptom), 1 (no plantar stepping in one hind leg), 2 (no plantar stepping in two hind legs or slight ankle movement in one hind leg), 3 (slight ankle movement in two hind legs or no ankle movement in one hind leg), 4 (no ankle movement in one hind leg and slight in the other one) to 5 (no ankle movement in both hind legs, almost loss of all movement ability), as described previously (26). The mice scored for 5 were euthanized as a humane endpoint according to the animal protocol.

### Real-time reverse transcription PCR

Animal tissues or whole blood (30μl) were collected in 330μl of lysis buffer (Invitrogen RNAasy mini-prep kit). The heart, liver, lung, brain and spleen were minced with a pair of scissors and homogenized in RLT buffer using an electrical pellet pestle (Kimble Chase LLC, USA). RNA was extracted following the Invitrogen RNAasy mini-prep kit protocol, and reverse-transcribed into cDNA using the TAKARA PrimeScriptTM RT Reagent Kit with gDNA Eraser (Perfect Real Time) (#RR047A). qPCR was performed with gene-specific primers and Bio-Rad SYBR Premix. qPCR results were calculated using the –ΔΔCt method and beta actin gene as an internal control. The qPCR primers are summarized in supplemental material Table S2.

### Plaque-forming assay

The viral particles in tissue homogenates or cell culture medium were quantitated by plaque forming assays as previously described (39). Briefly, samples were serially diluted by 10-fold using DMEM without FBS, and then 500 µL of each diluted sample plus 500 µL DMEM without FBS were added onto a Vero cell monolayer in a 6-well plate. The 6-well plate was then incubated at 37 °C for 2 h. The inoculum was replaced with 1% SeaPlaque agarose (Cat# 50100, Lonza) in complete DMEM medium (2 ml). The plate was left at room temperature for 30min to allow the agarose to solidify, and then moved to a cell culture incubator (37 °C, 5% CO_2_). Plaques were visualized by a Neutral Red exclusion assay after 3 days.

### Western blotting assay

Western blotting analysis was performed using standard procedures. Briefly, protein samples were resolved by SDS-PAGE (sodium dodecyl sulfate-polyacrylamide gel electrophoresis) and transferred to a nitrocellulose membrane. The membrane blot was incubated with a primary antibody over night at 4 °C, washed briefly and incubated with a HRP-conjugated secondary antibody for 1 hour at room temperature. An ultra-sensitive ECL substrate was used for detection (ThermoFisher, Cat# 34095).

### Co-immunoprecipitation

Trophoblasts or HEK293 cells were seeded at a density of 5×10^5^ cells/ml and transfected with protein expression plasmids (pVSVG, 1-80, 51-end, 1-end of FLAG-MSR1, and vector) using TransIT-X2® Transfection Reagent. Twenty-four hours after transfection, whole-cell extracts were prepared from transfected trophoblasts in lysis buffer (150 mM NaCl, 50 mM Tris pH 7.5, 1 mM EDTA, 0.5% NP40, 10% Glycerol) and were incubated with 50μl of anti-FLAG magnetic beads for 2hrs at 4°C. The whole procedure of co-immunoprecipitation was performed according to manufacturer’s protocol (Anti-Flag Magnetic Beads, Sigma-Aldrich). For co-immunoprecipitation of live VSV particles, 5×10^5^ cells/ml HEK293 cells were transfected with protein expression plasmids (1-80, 51-end, 1-end of FLAG-MSR1, and vector). Twenty-four hours later, cells were lysed in lysis buffer and incubated with 50μl of anti-FLAG magnetic beads for 2hrs at 4°C. The beads were washed three times with sterile PBS to completely remove detergent and unbound cellular proteins, and were then incubated with VSV-GFP virions (500μl, 1×10^7^ PFU/ml) for 2hrs at 4°C. The beads were washed with sterile PBS three times to remove unbound virions and the bound virions were eluted with 3xFLAG peptide in sterile PBS. The VSV-GFP pulled down by FLAG-MSR1 and fragments were then used to infect Vero cells or immunoblotting (Fig. S2b).

### Multiplex ELISA (enzyme-linked immunosorbent assay)

BioLegend’s LEGENDplex(tm) bead-based immunoassays of serum samples were performed exactly according to the instruction of BioLegend’s LEGENDplex(tm) kit. The flow cytometry data were analyzed using the LEGENDplex(tm) data analysis software.

### Graphing and statistics

The data from all experiments were analyzed using a Prism GraphPad Software and expressed as mean ± S.D. Survival data were analyzed using a Log-rank (Mantel-Cox) test. For *in vitro* cell culture results, a standard two-tailed unpaired Student’s t-test was used. For animal studies, an unpaired two-tailed nonparametric Mann-Whitney U test was applied to statistical analysis. *P* values ≤ 0.05 were considered significant. The statistic statements of experiments are detailed in each figure legend.

## Acknowledgements

This work was supported by a National Institutes of Health grant R01AI132526 and UConn Health startup fund to P.W.

## Author contributions

D.Y. designed and performed all the experimental procedures and data analyses. T.L., A.H., and T.G provided technical support and/or data analyses. H.W. assisted with data interpretation and discussion. P.W. conceived and oversaw the study. D.Y. and P.W. wrote the paper and all the authors reviewed and/or modified the manuscript.

