## Supplemental material and Figures for "A critical role for MSR1 in vesicular stomatitis virus infection of the central nervous system"

**Title page**

Running title: MSR1 is a cellular receptor for VSV

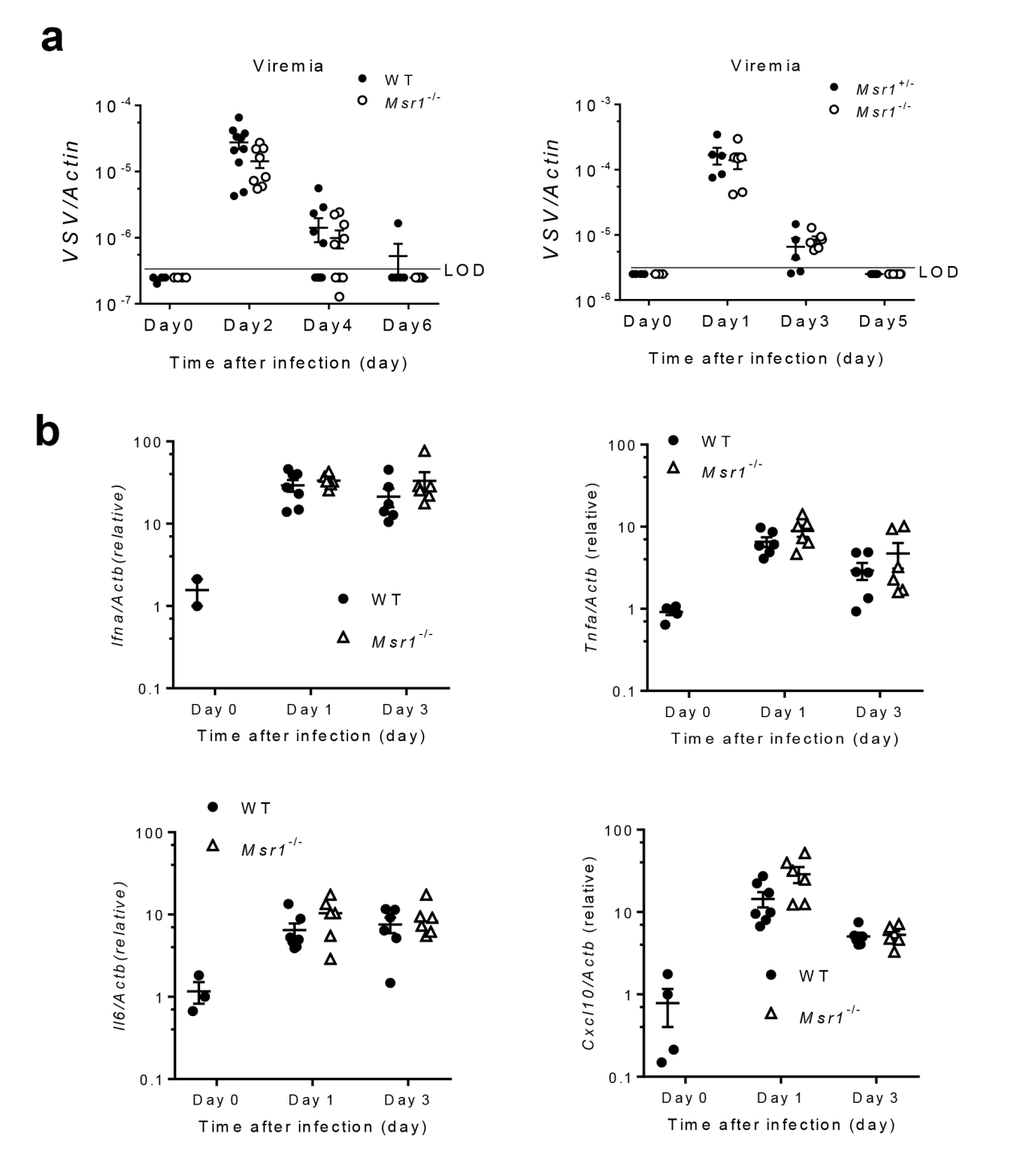

**Fig. S1 Msr1 is dispensable for systemic VSV dissemination and innate immune responses.** **a)** The viremia of WT mice, *Msr1^+/-^* and *Msr1^-/-^* littermates infected with 1×10^6^ PFU/mouse of VSV-GFP by retro-orbital injection, assessed by quantitative RT-PCR, N=8-9 mice/group for WT and *Msr1^-/-^*, N=5 mice/genotype for *Msr1^+/-^* and *Msr1^-/-^* littermates. **b)** The mRNA levels of *Ifnα*, *Tnfα*, *Il6* and *Cxcl10* in leukocytes of WT and *Msr1^-/-^* mice after VSV infection, assessed by quantitative RT-PCR, N=6 mice/group. All the data are presented as mean ± S.E.M. and statistical significance are analyzed by non-parametric Mann-Whitney U test.

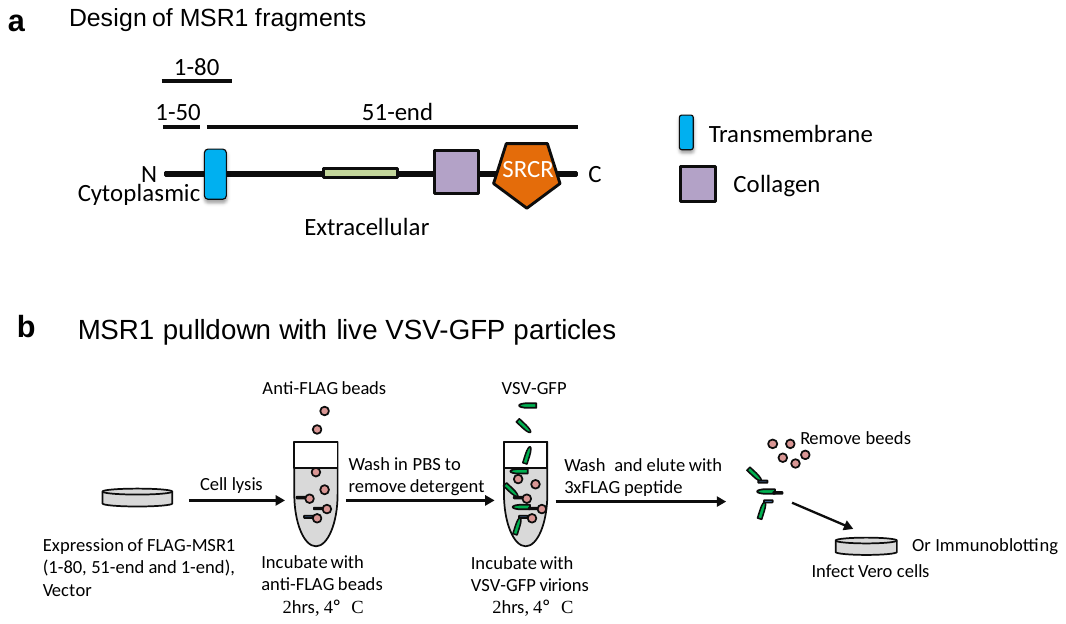

**Fig. S2 Design of MSR1 fragments and MSR1 pulldown with live VSV-GFP virions. a)** Amino residues 1-50: cytoplasmic N-tail, 1-80: cytoplasmic N-tail plus transmembrane, and 51-end: extracellular domains including the collagen, scavenger receptor cysteine-rich (SRCR) and the transmembrane domains. b) Procedure of MSR1 pulldown with live VSV-GFP particles using FLAG-MSR1 and its fragments.

Supplemental Table S1

**Primers for gene subcloning and expression with restriction enzyme sites**

| Gene | Species | Forward primer (5’-3’) | Reverse primer (5’-3’) | Enzyme sites |
| --- | --- | --- | --- | --- |
| *MSR1* | Hs | CTTGCGGCCGCGGAGCAGTGGGATCACTTTCA | TTGGATCCCTATAAAGTGCAAGTGACTCCAGC | NotI and BamHI |
| *MSR1* fragment *1-50* | Hs | CTTGCGGCCGCGGAGCAGTGGGATCACTTTCA | TTGGATCCCTAAGCTTTGAAGGACTTCAGTT | NotI and BamHI |
| *MSR1* fragment *1-80* | Hs | CTTGCGGCCGCGGAGCAGTGGGATCACTTTCA | TTGGATCCCTACGTTTCCCACTTCAGGAG | NotI and BamHI |
| *MSR1* fragment *51-end* | Hs | CTTGCGCCCGCGGCACTGATTGCCCTTTAC | TTGGATCCCTATAAAGTGCAAGTGACTCCAGC | NotI and BamHI |

Supplemental Table S2

**Primers for qPCR**

| Gene | Species | Forward primer (5’-3’) | Reverse primer (5’-3’) |
| --- | --- | --- | --- |
| VSV | Indiana strain | TGATACAGTACAATTATTTTGGGAC | GAGACTTTCTGTTACGGGATCTGG |
| Ldlr | Mm | GAATCTACTGGTCCGACCTGTC | CTGTCCAGTAGATGTTGCGGTG |
| Msr1 (Scara1) | Mm | AGTGCTGTCTTCTTTACCAGC | GTGAGGAAGGGATGCTGTA |
| Scara2 | Mm | ATGGCACCAAGGGAGACAAAGG | GCCTGGTTTTCCAGCATCACCT |
| Scara3 | Mm | CCACGGAGAAATCCTTCGCAATG | TAGGTCCTCTGCTACCAACAGG |
| Scara4 | Mm | ACTCCAAGCACGGTCAGCTCAT | CTTGTTGCCAGTTGGACCAGGT |
| Scara5 | Mm | TTTGATGGCAGGAGCCTGTCCA | CCCACAAGAATCAGGAAGACCAG |
| Scarb1 | Mm | ACACCCGAATCCTCGCTGGAAT | CCGTTGGCAAACAGAGTATCGG |
| Scarb2 | Mm | TAGCCAACACCTCCGAAAACGC | CGAACTTCTCGTCGGCTTGGTA |
| Scarb3 (CD36) | Mm | GGACATTGAGATTCTTTTCCTCTG | CAAAGGCATTGGCTGGAAGAAC |
| Msr2 | Mm | CCTGATCCAGAGTGCAATCGTG | CACATCTCCGATGAAGGGCAAG |
| Actin | Mm | AGAGGGAAATCGTGCGTGAC | CAATAGTGATGATGACCTGGCCGT |
| Ifna | Mm | CTTCCACAGGATCACTGTGTACCT | TTCTGCTCTGACCACCTCCC |
| Tnfa | Mm | CTCCAGGCGGTGCCTATGT | GAAGAGCGTGGTGGCCC |
| Il6 | Mm | CCAGAAACCGCTATGAAGTTCC | TCACCAGCATCAGTCCCAAG |
| Cxcl10 | Mm | ATCATCCCTGCGAGCCTATCCT | GACCTTTTTTGGCTAAACGCTTTC |
